# The ASD Risk Gene *D5Ertd579e* Regulates Synaptic Plasticity and Selective Autism-Related Behaviors

**DOI:** 10.64898/2026.06.02.729632

**Authors:** Isidora Stankovic, Pablo Lituma, Eva Onur, Megan Nguyen, Dilan Rasool, Dilek Colak

**Affiliations:** Center for Neurogenetics, Feil Family Brain and Mind Research Institute, Weill Cornell Medical College, Cornell University, New York City, New York, 10021, USA; Gale & Ira Drukier Institute of Children’s Health, Weill Cornell Medical College, Cornell University, New York City, New York, 10021, USA

**Keywords:** Autism spectrum disorder, KIAA0232, D5Ertd579e, Synaptic plasticity, Neurodevelopment

## Abstract

Autism spectrum disorder (ASD) is a heterogeneous neurodevelopmental condition shaped by contributions from hundreds of genes, many of which remain poorly characterized. This largely uncharacterized genomic landscape may therefore hold critical insight into how diverse molecular disruptions converge on shared social phenotypes. Here, we investigated *KIAA0232* (mouse orthologue *D5Ertd579e*), an uncharacterized locus lacking known functional domains, using a global null knockout mouse model. While loss of *D5Ertd579e* did not overtly disrupt cortical progenitor dynamics, laminar organization, or gross brain morphology, *D5Ertd579e* null mutants exhibited selective behavioral deficits in vocalization, sociability, and novelty preference, while anxiety- and memory-related behaviors remained preserved. These behavioral phenotypes were accompanied by attenuated long-term plasticity, despite normal basal synaptic transmission. Together, our findings indicate that *D5Ertd579e* loss selectively alters neurodevelopment, preferentially impacting neural systems involved in social and motivational processing while preserving hippocampal-dependent networks. We propose that *D5Ertd579e* functions as a regionally specific regulator of neurodevelopment, whose disruption may contribute to ASD through distinct genetic pathways. More broadly, this study underscores the importance of interrogating uncharacterized loci to refine mechanistic models of the social brain in ASD.

## Introduction

Autism spectrum disorder (ASD) is a heterogeneous neurodevelopmental condition characterized by alterations in social communication, restricted or repetitive behaviors, and atypical sensory processing. Affecting approximately 1 in 32 children in the United States (Shaw et al., 2025), ASD presents a clinical and societal challenge due to its phenotypic variability and the absence of curative interventions. From a biological standpoint, ASD etiology is highly complex, involving contributions from hundreds of genes, epigenetic regulators, and environmental factors. Despite substantial progress in gene discovery, much of the genetic architecture underlying ASD remains poorly understood, with many implicated loci lacking functional annotation.

Genetic studies have repeatedly identified mutations in genes such as SHANK3, NRXN1, NLGN3, and CHD8, all of which play important roles in synaptic signaling, transcriptional regulation, and chromatin remodeling (Bourgeron, 2009; De Rubeis et al., 2014). These findings support theoretical models including the synaptopathy hypothesis and the excitation–inhibition imbalance model, both of which emphasize disrupted neuronal connectivity as a central feature of ASD pathology (Rubenstein & Merzenich, 2003; Nelson & Valakh, 2015). But this well-characterized subset of genes represents only a fraction of the ASD-associated genomic landscape. A substantial proportion of implicated loci remain unclassified, lacking defined molecular functions or structural domains, and are therefore underrepresented in mechanistic studies. However, whether uncharacterized ASD-associated genes contribute to selective circuit-level and behavioral phenotypes remains unclear.

One of these overlooked genes is KIAA0232 (mouse orthologue *D5Ertd579e*), an uncharacterized locus with no recognizable functional domains or family associations in current databases. Publicly available annotations from the Mouse Genome Informatics database suggest that disruption of *D5Ertd579e* is associated with behavioral and neurodevelopmental phenotypes, although its functional role remains largely uncharacterized. Despite suggestive phenotypic annotations, KIAA0232 has not been functionally characterized in the context of brain development or ASD. *D5Ertd579e* is expressed across multiple stages of mouse brain development and is detected in diverse neural cell populations, with conserved expression observed in human brain tissue (Di Bella et al., 2021; Loo et al., 2019). Large-scale sequencing studies have demonstrated that de novo coding mutations contribute significantly to ASD risk, particularly in genes involved in neurodevelopmental and synaptic processes (Iossifov et al., 2014; Satterstrom et al., 2020). Subsequent studies integrating rare coding variants and copy number variants have expanded the ASD risk gene landscape and identified additional moderate-risk genes with developmental and neuronal functional relevance (Zhou et al., 2022; Fu et al., 2022). However, the functional contribution of such unannotated genes to neurodevelopment and behavior remains largely unknown.

Emerging evidence suggests that ASD-related phenotypes often arise from disruptions in specific neural circuits rather than global brain dysfunction. In particular, corticostriatal and cerebellar networks have been increasingly implicated in social behavior, motivation, and motor coordination, complementing classical models centered on limbic circuitry (Fucillo, 2016). At the same time, alterations in developmental timing and critical period regulation are recognized as key features of ASD, with disruptions in processes such as cortical organization and synaptic refinement contributing to long-term circuit dysfunction (LeBlanc & Fagiolini, 2011; Schafer et al., 2012). Together, these frameworks suggest that genes influencing neurodevelopment may exert selective effects on discrete circuits and behavioral domains.

Here, we investigate the role of *D5Ertd579e* in cortical development and behavior using a full knockout mouse model, with a focus on whether disruption of an uncharacterized gene produces selective neurodevelopmental and behavioral phenotypes. We demonstrate that loss of *D5Ertd579e* does not overtly disrupt cortical progenitor dynamics, laminar organization, or gross brain morphology, but instead produces selective behavioral alterations in vocalization, sociability, and novelty preference, with preserved anxiety- and memory-related functions. These findings suggest that *D5Ertd579e* contributes to neurodevelopment in a circuit-selective manner, preferentially affecting pathways involved in social and motivational processing.

More broadly, our work highlights the importance of interrogating uncharacterized genes to refine mechanistic models of ASD. By linking *D5Ertd579e* disruption to selective behavioral phenotypes in the absence of gross anatomical abnormalities, we provide evidence that subtle developmental perturbations in specific neural circuits may underlie core features of ASD. This supports a model in which diverse genetic risk factors converge on shared behavioral outcomes through regionally and temporally specific effects on brain development.

## Methods

### *D5Ertd579e* knockout mouse

*D5Ertd579e* knockout mice were obtained from the Jackson Laboratory (MMRRC Strain #043784-JAX) and were generated by CRISPR/Cas9 genome editing (Adams, D.J., Barlas, B., McIntyre, R.E. *et al*., 2024). A 242 bp segment was deleted, eliminating exon 4 together with 175 bp of adjacent intronic sequence that included both splice donor and acceptor sites. The resulting frameshift is predicted to alter the coding sequence after amino acid 122 and introduce a premature stop codon.

### Immunohistochemistry

Mice underwent transcardial perfusion with 4% paraformaldehyde (following a PBS flush). Brains were post-fixed overnight, transferred into 30% sucrose, and stored at 4 °C. Tissue was embedded in OCT compound and snap-frozen at −80 °C for coronal cryosectioning at either 30 μm or 300 μm thickness. Sections were permeabilized in 1% Triton X-100 for 30 min and subsequently blocked in 10% normal goat serum for 1 h. Primary antibody incubation was performed overnight at 4 °C. On the following day, sections were incubated with secondary antibodies for 2 h at room temperature. Nuclear counterstaining with Hoechst or DAPI was included in the final wash before mounting. Primary antibodies: DAPI (ThermoFisher; D1306), SOX2 (R&D Systems; MAB2018-SP), KIAA0232 (ThermoFisher; PA5-57862), TBR2 (EMD Millipore; AB15894), CTIP2 (Abcam; ab18465), CUX1 (ThermoFisher; 11733-1-AP), Hoechst (ThermoFisher; H3570), Gad67 (ThermoFisher; MA5-24909), NeuN (Invitrogen; PA5-143567), Syt2 (Osenses, OSS00023W), and Gephyrin (Invitrogen; MA5-25368). Secondary antibodies: Alexa 488 (anti-rabbit; ThermoFisher, A-11008) and 647 (anti-mouse; ThermoFisher, A-21235 and anti-chicken; ThermoFisher, A-21449).

### Quantitative RT-PCR

Total RNA was extracted using the Zymo RNA extraction kit (R2053) according to the manufacturer’s protocol. From this, 500 ng of RNA was reverse transcribed into cDNA with oligo(dT) and random primers using the cDNA synthesis kit (ThermoFisher, AB1453B).

Quantitative real-time PCR (qRT-PCR) was carried out with the SsoAdvanced™ Universal SYBR® Green Supermix (Bio-Rad, 1725270), using 10 ng of cDNA per reaction. Primer pair efficiencies were assessed by performing qPCR on tenfold serial dilutions of cDNA and calculating linear regression of Ct values within the 20–30 cycle range. Once primer validity was confirmed, mRNA quantification was carried out using the ΔΔCt method (Colak et al., 2023).

Relative transcript levels were normalized to ACTB mRNA.

### Behavioral assays

For all behavioral assays, mice were habituated to the testing room for at least 30 min, and experiments were conducted at the same time of day unless otherwise noted. Context- and cue-dependent fear conditioning was used to evaluate Pavlovian associative learning and fear learning. In this paradigm, mice learn to associate an auditory tone (conditioned stimulus, CS) with an aversive foot shock (unconditioned stimulus, US; 1 s/0.9 mA, delivered five times).

Freezing behavior was measured as the conditioned response (CR) and served as the primary readout of learning (day 1) and memory retrieval (days 2–3). On day 1, after a 2 min habituation period in the conditioning chamber, each mouse received five consecutive 30 s tone presentations (CS), each co-terminating with a foot shock (US). Learning was evaluated by plotting the percentage of freezing across successive tone presentations. On day 2, mice were re-exposed to the original conditioning context for 6 min without tone or shock to test context-dependent associative memory. On day 3, retrieval of the tone-shock association was assessed by placing animals in a novel context where only the CS was presented. The freezing percentage was averaged across all CS presentations as a measure of tone-dependent fear learning. On all days, freezing behavior was quantified using automated software (Med Associates Inc.).

Additional behavioral assays to assess locomotor activity, anxiety-like behavior, and repetitive behaviors were conducted as described previously (Colak et al., 2023).

For ultrasonic vocalization (USV) analysis, pup calls were recorded at postnatal day 8 (P8). Pups were briefly separated for 20-30 minutes from the dam and placed in a clean container maintained at room temperature prior to testing. Vocalizations were recorded for 3 minutes using an ultrasonic microphone and specialized acquisition software (Aviosoft Recorder), and call number, duration, and frequency parameters were quantified to assess communication behaviors.

### Hippocampal slice preparation

Male and female mice were anesthetized with 4% isoflurane and perfused with 25 mL of cold NMDG solution containing (in mM): 93 NMDG, 2.5 KCl, 1.25 NaH□PO□, 30 NaHCO□, 20 HEPES, 25 glucose, 5 sodium ascorbate, 2 thiourea, 3 sodium pyruvate, 10 MgCl□, and 0.5 CaCl□, adjusted to pH 7.35 with HCl (SA49, Fisher Scientific). Following perfusion, brains were extracted, and hippocampi were dissected and sectioned in cold NMDG solution using a VT1200S microslicer (Leica Co.).

Acute hippocampal slices (300–400 µm) were collected and transferred to a chamber containing artificial cerebrospinal fluid (ACSF) composed of (in mM): 124 NaCl, 2.5 KCl, 26 NaHCO□, 1 NaH□PO□, 2.5 CaCl□, 1.3 MgSO□, and 10 glucose. The slice chamber was maintained at 33– 34°C in a water bath. Ten minutes after completing slice collection, the chamber was brought to room temperature, and slices were allowed to recover for at least 45 minutes before experimentation. All solutions were continuously oxygenated with 95% O□ and 5% CO□ and maintained at pH 7.4.

### Spine Density Assessment

Animals were monitored for 72 hours following stereotaxic surgery. Approximately 10 days later, mice were anesthetized with 4% isoflurane and intracardially perfused with PBS to remove blood, followed by 4% paraformaldehyde (PFA) for fixation. Brains were post-fixed overnight in 4% PFA and subsequently cryoprotected in 30% sucrose. Once cryoprotection was complete, brains were embedded in tissue trays, frozen in OCT compound, and stored at –80°C. Coronal hippocampal sections (250–300 µm) were obtained using a Leica CM3050S cryostat, mounted onto microscope slides, and stored at –20°C until further processing.

The following day, coronal sections were immunostained to enhance EGFP fluorescence using a chicken anti-GFP antibody (1:50, A10262, Thermo Fisher Scientific), following previously described immunostaining procedures. Slides were air-dried in the dark and ProLong Antifade Glass Mountant was applied for coverslip adhesion.

Fluorescent imaging was performed using an Olympus Fluoview FV1000 confocal microscope at 640 × 640-pixel resolution with the 488 nm laser line and a 60× oil-immersion objective (1.4 NA, Nikon) at 4.0 µs/pixel and 8× digital zoom. Z-stack images (10–15 µm thickness; 0.2 µm step size) were collected from dendritic regions of CA1 pyramidal neurons, at least 150 µm from the soma. Images were processed in Fiji to generate Z-projections from at least five fields of view per sample/genotype. Spine density was quantified in 10 µm dendritic segments in a blinded manner for both wild-type and knockout conditions.

### Electrophysiology

Electrophysiological recordings were performed at 31.2 ± 1 °C under temperature control using a TC-344C system (Warner Instruments) in a submersion-type recording chamber continuously perfused with ACSF at 2 mL/min. A stimulating glass electrode filled with ACSF was positioned in the stratum radiatum to activate Schaffer collateral inputs via an Isoflex stimulus isolator (A.M.P.I.) delivering pulses of 100 µs duration. Extracellular field excitatory postsynaptic potentials (fEPSPs) were recorded using patch-type pipettes filled with 1 M NaCl.

For long-term potentiation (LTP) experiments, 400 µm-thick slices were stimulated every 20 s for a 15 min baseline period. LTP was induced by theta-burst stimulation (TBS) consisting of 10 bursts of 5 pulses at 100 Hz, delivered every 200 ms (inter-burst interval), and repeated four times at 5 s intervals, as previously described (104). LTP recordings were performed in the presence of picrotoxin (100 µM; P1675, Sigma-Aldrich) and CGP-55845 (3 µM; No. 1248, Tocris Bioscience) to inhibit synaptic transmission. The magnitude of potentiation was quantified by comparing the average fEPSP slope during the last 10 min of recording with the baseline average.

For input–output and paired-pulse ratio (PPR) experiments, synaptic inputs were stimulated every 10 s. Stimulation intensity (Isoflex stimulator) was increased from 0–20 V in 5 V increments. fEPSP slopes and fiber volley amplitudes were analyzed using ClampFit 11.2 software. PPR was assessed by delivering two stimuli at inter-stimulus intervals ranging from 10–500 ms and calculated as the ratio of fEPSP slopes (fEPSP□/fEPSP□). Both input–output and PPR recordings were conducted in the absence of pharmacological agents.

Field recordings were amplified using a MultiClamp 700B amplifier and Axon Digitizer 1550B (Molecular Devices), filtered at 2 kHz, and digitized at 5 kHz. Stimulation and data acquisition were controlled with Clampex 11.2 software.

### Statistical Analysis

Statistical analyses and data visualization were conducted using GraphPad Prism 10. Data distributions were assessed for normality with the Shapiro–Wilk test. Parametric unpaired *t*-tests were applied to normally distributed data, while non-parametric Mann–Whitney tests were used for skewed distributions. For behavioral experiments, Sidak’s multiple comparison test was used to analyze Day 1 of the contextual fear conditioning paradigm.

## RESULTS

### *D5Ertd579e* knockout validation and gross morphological analysis

To validate the *D5Ertd579e* knockout model, we first quantified transcript and protein expression across developmental stages. Publicly available transcriptomic datasets indicate that *D5Ertd579e* is expressed throughout multiple stages of mouse brain development and across diverse neural cell populations, including neuronal and progenitor-like populations, among others (Di Bella et al., 2021; Loo et al., 2019). qRT-PCR analysis of adult cortical tissue revealed a marked reduction (∼80%) in *D5Ertd579e* mRNA levels in knockout (KO) animals relative to wild-type (WT) littermates (**Fig. 1A**). Consistent with this, immunostaining of embryonic brain sections demonstrated a strong reduction in D5Ertd579e protein signal in KO mice (**Fig. 1E**), confirming effective gene disruption at both transcript and protein levels.

**Figure 1.**
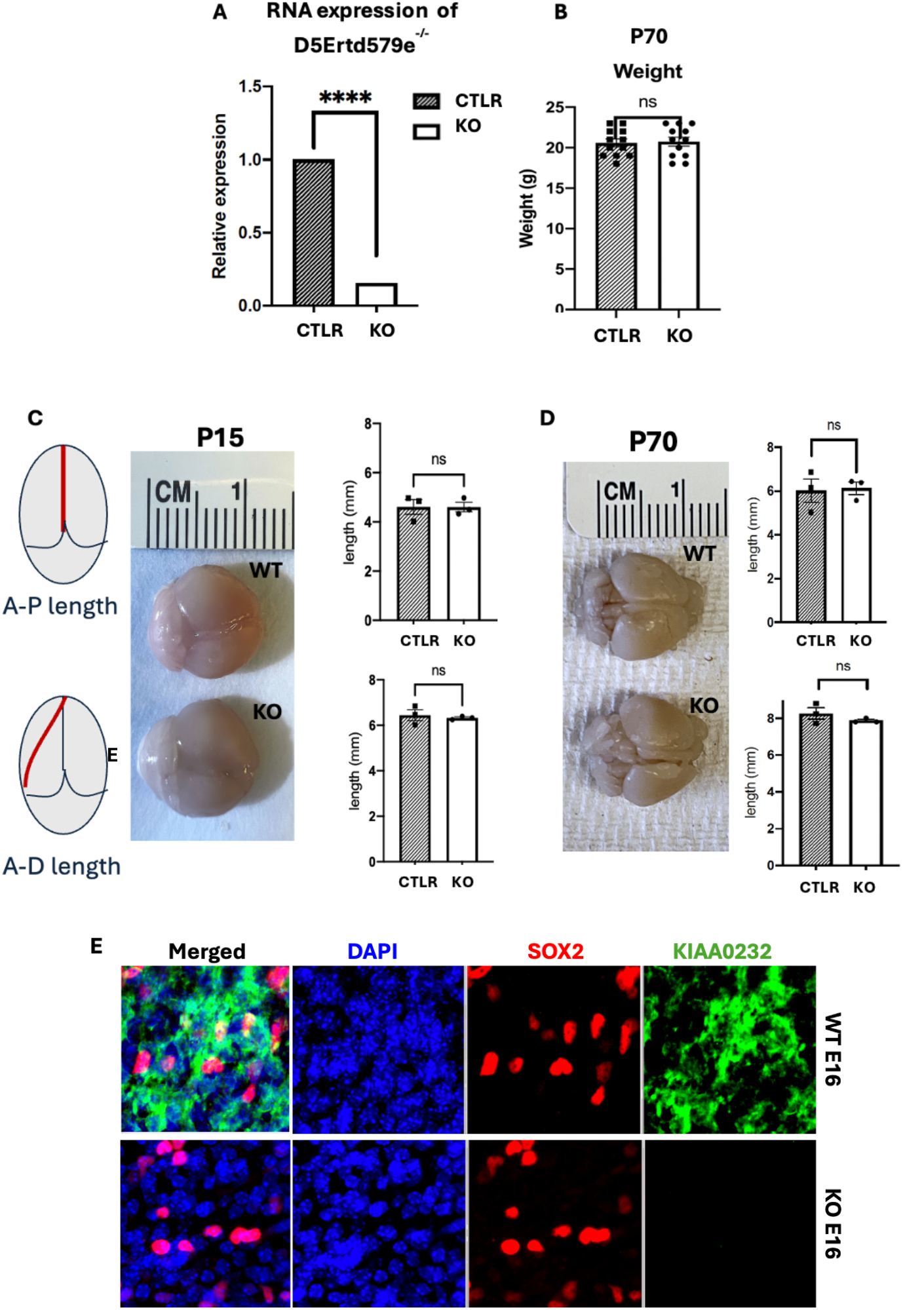
Characterization and validation of *D5Ertd579e* knockout mice. (A) RT-qPCR confirms a marked reduction of *D5Ertd579e* transcript in knockout (KO) cortex compared with WT littermates. (B) Body weight at postnatal day 70 (P70) shows no significant difference between wild-type (WT, n = 12) and knockout (KO, n = 12) mice. (C) Anterior-posterior and anterior-dorsal brain length at P15 is unchanged between groups (WT n = 3, KO n = 3), representative image shown. (D) Likewise, at P70, measurements reveal no significant differences between groups (WT n = 3, KO n = 3). (E) D5Ertd579e protein localizes to both nuclear and cytoplasmic compartments. Data are presented as mean ± SEM. *n* represents biological replicates. *ns* represents not significant. Unpaired student t-test was employed. *p* < 0.05.

We next assessed whether *D5Ertd579e* deletion impacts overall growth or brain morphology. Body weight measurements at postnatal day 70 (P70) showed no significant difference between KO and WT animals (**Fig. 1B**). Similarly, quantitative analysis of brain dimensions at both early postnatal (P15) and adult (P70) stages revealed no differences in anterior–posterior, dorsal– ventral, or medial–lateral axes (**Fig. 1C-D**). Together, these findings indicate that *D5Ertd579e* is not required for gross somatic growth or global brain development.

### *D5Ertd579e* knockout produces selective social and behavioral alterations

To determine whether loss of *D5Ertd579e* affects behavior, we performed a wide array of assays assessing communication, repetitive behavior, locomotion, and social interaction. Early-life social communication was assessed using ultrasonic vocalization (USV) assays. At postnatal day 8 (P8), KO pups emitted significantly fewer vocalizations in response to maternal separation compared to WT controls (**Fig. 2A**), indicating impaired early social communication.

**Figure 2.**
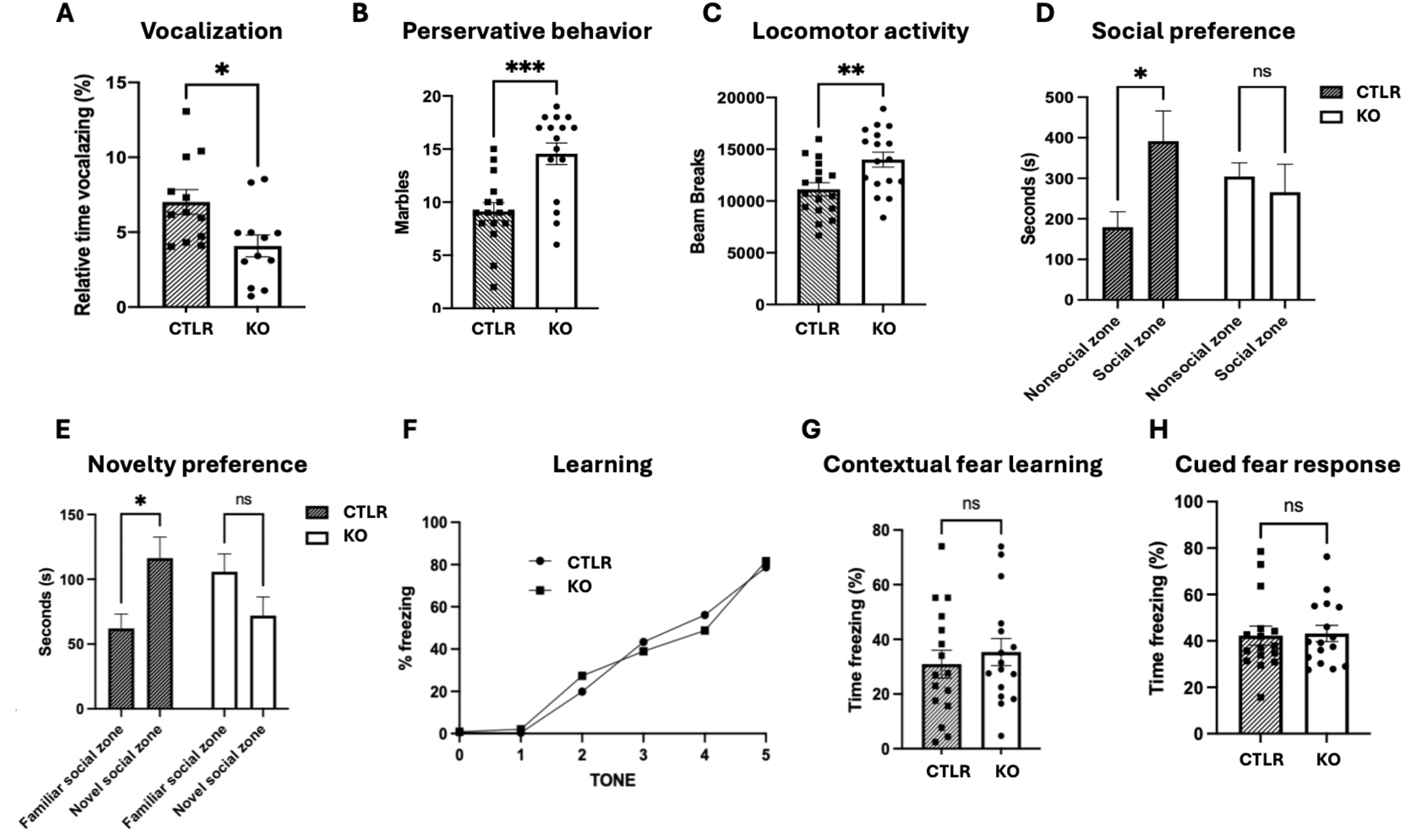
*D5Ertd579e* knockout mice exhibit selective social and behavioral alterations. (A) Ultrasonic vocalization deficits: *D5Ertd579e* knockout mice display significantly reduced relative time spent vocalizing during early postnatal development compared to wild-type controls (WT n = 16, KO n = 16; *p < 0.05, unpaired t-test). (B) Enhanced perseverative behavior: KO mice bury significantly more marbles in the marble-burying assay, reflecting increased repetitive/perseverative tendencies (WT n = 16, KO n = 16; ***p < 0.001, unpaired t-test). (C) Locomotor hyperactivity: KO mice show increased locomotor activity measured as total beam breaks in open-field testing (WT n = 16, KO n = 16; **p < 0.01, unpaired t-test). (D-E) Impaired social and novelty preference: WT mice demonstrate significant preference for the social zone over the non-social zone in three-chamber testing, while KO mice show no preference between zones (WT social preference: *p < 0.05; KO preference: ns, two-way ANOVA with post-hoc comparisons). (F) KO and WT mice showed comparable acquisition of tone–shock association during the training phase, with both groups increasing freezing across trials. (G) In the contextual fear test, KO mice froze at similar levels to WT littermates, indicating intact hippocampal-dependent memory. (H) In the cued fear test, KO mice also exhibited freezing responses comparable to WT, suggesting normal amygdala-dependent associative learning. Data are presented as mean ± SEM; ns denotes non-significance.

Repetitive and exploratory behaviors were also altered. In the marble-burying assay, KO mice buried significantly more marbles than WT mice (**Fig. 2B**), consistent with increased perseverative or repetitive behavior. In parallel, open-field testing revealed a significant increase in total locomotor activity in KO mice, as measured by beam breaks over the testing period (**Fig. 2C**), suggesting heightened exploratory or hyperactive behavior.

Social interaction assays revealed a selective deficit in sociability. In the three-chamber test, WT mice showed a clear preference for interacting with a novel conspecific over an empty chamber, whereas KO mice failed to exhibit this preference (**Fig. 2D**). Furthermore, KO mice did not display a preference for a novel over a familiar conspecific (**Fig. 2E**), indicating impaired social novelty recognition. Importantly, these deficits were not attributable to reduced general activity, as KO mice exhibited increased locomotion in the open field. Together, these findings demonstrate that *D5Ertd579e* deletion selectively disrupts social communication and interaction while enhancing repetitive and exploratory behaviors.

### *D5Ertd579e* disruption does not cause learning and memory deficits in mice

To assess whether *D5Ertd579e* deletion leads to broader cognitive deficits, we evaluated associative learning and memory using fear conditioning paradigms. During the acquisition phase, both WT and KO mice exhibited comparable increases in freezing behavior across training trials, indicating intact learning of the tone–shock association. In the contextual fear test, KO mice displayed freezing levels similar to WT controls (**Fig. 2F-G**), suggesting preserved hippocampal-dependent memory. Likewise, cued fear conditioning, which depends on amygdala function, was also unaffected (**Fig. 2H**). These findings indicate that *D5Ertd579e* deletion does not impair core learning and memory processes, supporting a model in which behavioral alterations are domain-specific rather than global.

### Cortical progenitor dynamics and laminar organization are preserved following *D5Ertd579e* loss

Given the observed behavioral phenotypes, we next investigated whether *D5Ertd579e* deletion disrupts cortical development. D5Ertd579e expression was detected throughout cortical development, with signals present across cortical layers at embryonic stages E14–E18 (**Fig. 3A**). However, analysis of progenitor zones revealed no structural abnormalities. Immunostaining for SOX2 and TBR2 showed comparable organization and distribution of neural stem and intermediate progenitor cells within the ventricular zone (VZ) and subventricular zone (SVZ) in WT and KO embryos (**Fig. 3B-C**). Quantification of VZ thickness further confirmed no significant differences between genotypes.

**Figure 3.**
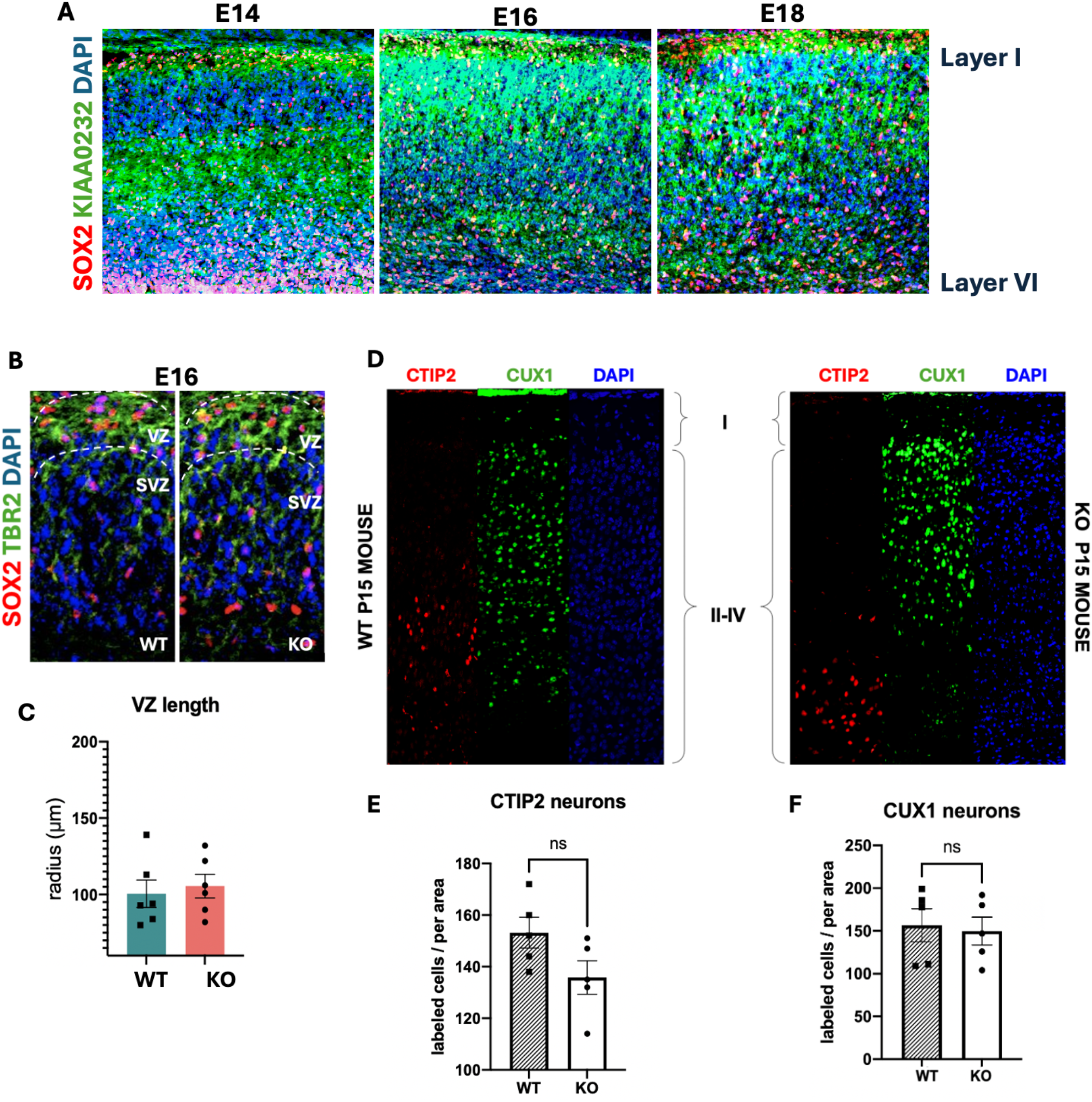
Cortical analyses in *D5Ertd579e* knockout brains. (A) Embryonic mouse brains collected and stained at embryonic day (E) 14, 16, and 18, show KIAA0232 expression in layer I-VI. (B) Representative confocal microscopy images of embryonic (E16) cerebral cortex from wild-type (WT, left panel) and D5Ertd579e knockout (KO, right panel) shown. Sections are immunolabeled for stem cell, SOX2, and progenitor cells, TBR2. Nuclei were stained with DAPI. Ventricular zone (VZ) and subventricular zone (SVZ) are delineated by dashed lines. (C) The radius of the ventricular zone showed no difference between WT and KO mice. (D) Representative coronal sections of the cerebral cortex from WT and KO mice at postnatal day 15 is shown. Cortical layers were stained with cortical layer markers CTIP2 and CUX1. DAPI was used as a nuclei marker. (E) CTIP2 neurons, and CUX1 (F) neurons were quantified in P15 mice, and showed no significant difference between groups. Each individual point represents a biological replicate; ns is non-significant; unpaired student t-test used.

Postnatal analysis at P15 demonstrated preserved cortical lamination. Layer-specific markers CTIP2 (layer V) and CUX1 (layers II–IV) revealed appropriate neuronal positioning and distribution in both WT and KO mice (**Fig. 3D-F**), with no significant differences in cell number or layer organization. Together, these findings indicate that D5Ertd579e is not required for the establishment of cortical progenitor architecture or laminar patterning.

### Loss of *D5Ertd579e* does not alter hippocampal inhibitory neuron numbers or synaptic density

To determine whether inhibitory circuitry is altered, we quantified GABAergic interneurons and inhibitory synapse density in the hippocampal CA1 region. Immunostaining for GAD67 revealed comparable numbers of inhibitory interneurons in WT and KO mice (**Fig. 4A-B**). In addition, analysis of parvalbumin-positive (PV□) somatic inhibitory synapses, identified by colocalization of Syt2 and Gephyrin puncta, showed no significant difference in synapse density between genotypes (**Fig. 4C-D**). These data indicate that *D5Ertd579e* loss does not disrupt inhibitory neuron number or the structural organization of inhibitory synapses in CA1.

**Figure 4.**
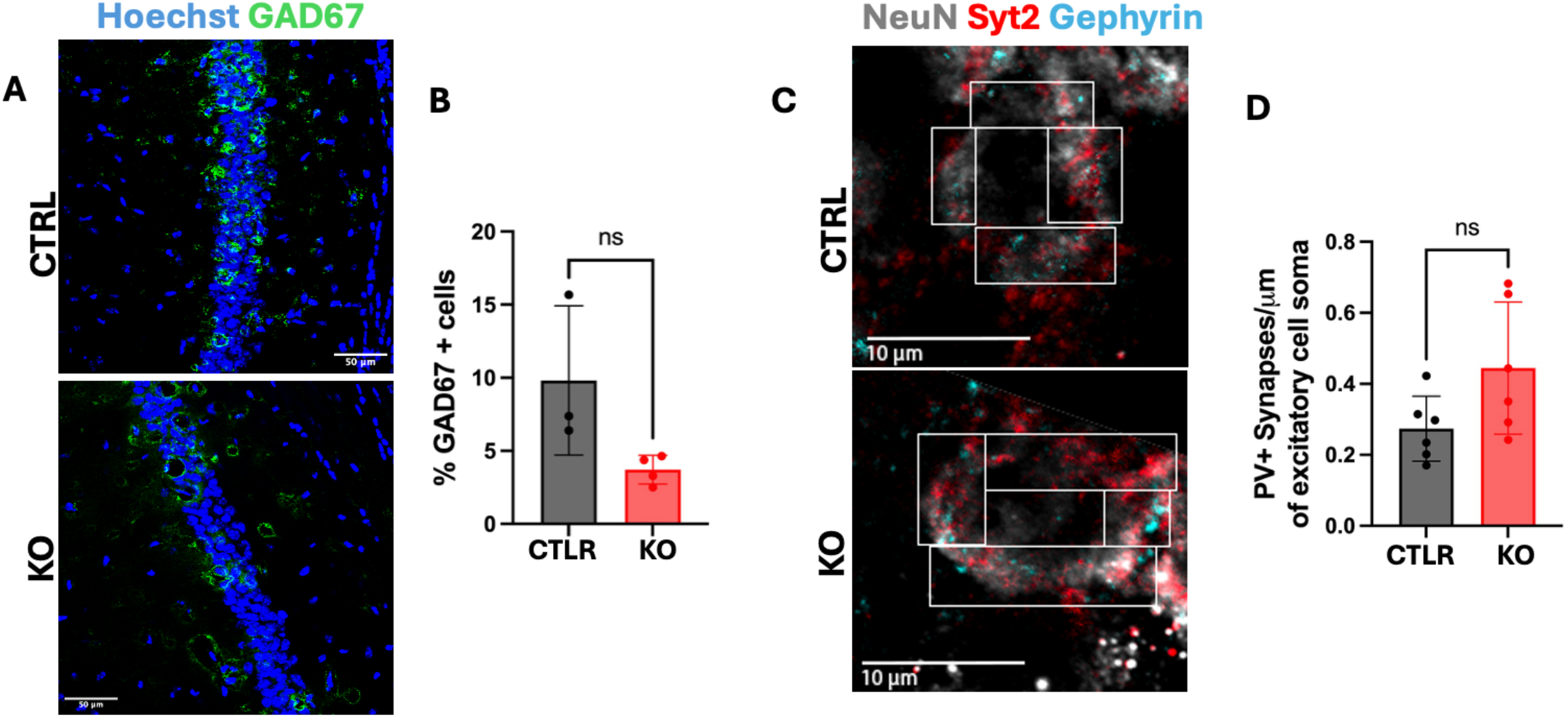
GABAergic cell number and somatic parvalbumin inhibitory synapse density is normal in the hippocampus of *D5Ertd579e* KO animals. (**A-B**) GAD67+ cell number in hippocampus CA1 area of P45 KO mice is unchanged as compared to CTRL (n = 3 mice per genotype, Mann-Whitney test, ns p > 0.05). (**C-D**) Parvalbumin somatic inhibitory synapse density of CA1 pyramidal neurons in P45 KO mice is similar to CTRL (n = 2 mice per genotype, Mann-Whitney test, ns p > 0.05).

### *D5Ertd579e* loss selectively impairs synaptic plasticity while preserving basal synaptic function

To further assess synaptic function, we examined both structural and electrophysiological properties of hippocampal CA1 neurons. Analysis of dendritic spine density revealed no significant difference between WT and KO mice (**Fig. 5A-B**), indicating that excitatory synapse density is preserved. Consistent with this, basal synaptic transmission was not significantly different, as demonstrated by similar input–output relationships between groups (**Fig. 5D-E**).

**Figure 5.**
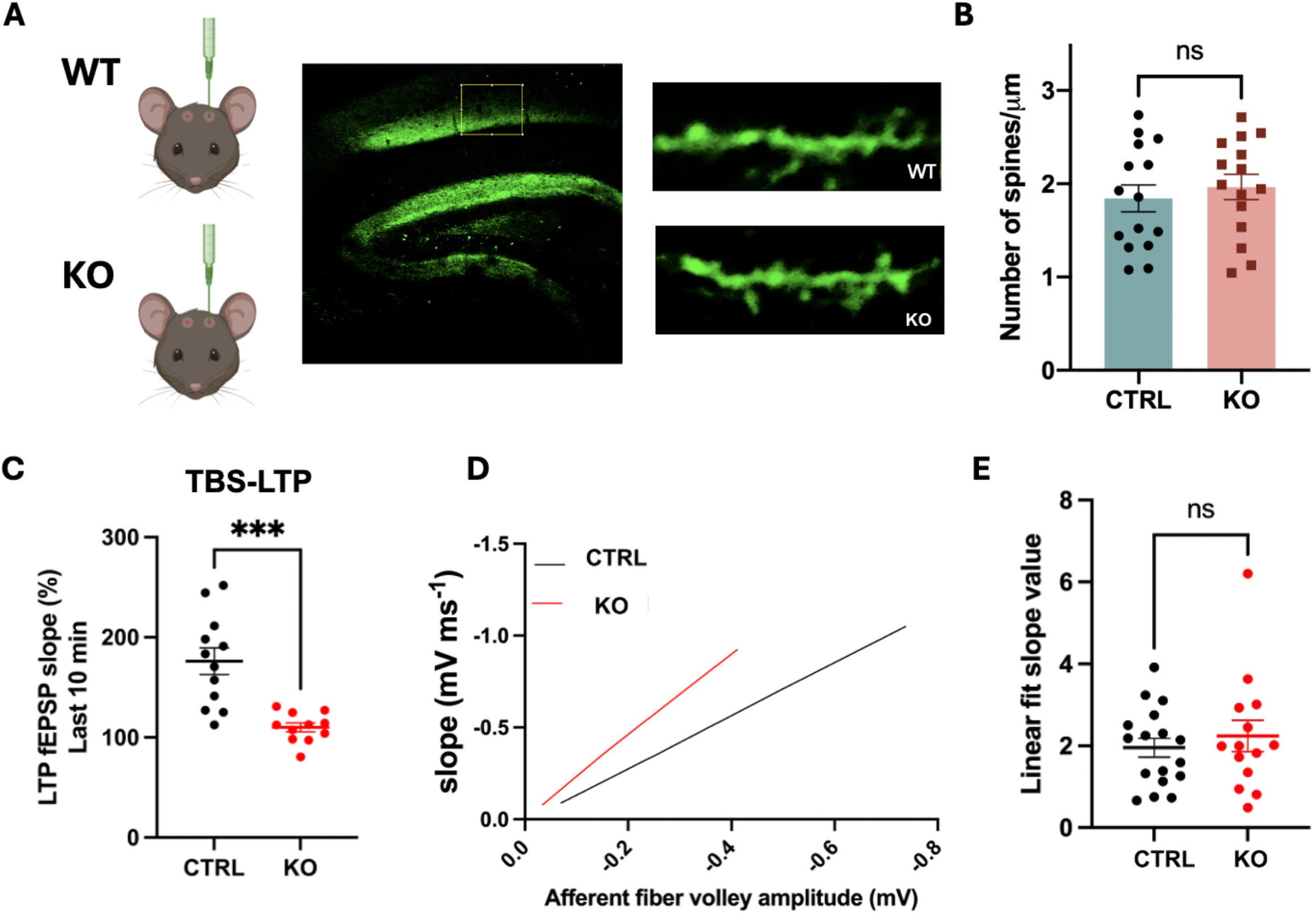
*D5Ertd579e* KO mice display normal hippocampal basal synaptic transmission but impaired long-term potentiation (LTP). (A) Representative schematic of viral injection and hippocampal CA1 region used for dendritic spine imaging. Example confocal images of apical dendrites from WT and KO mice are shown. (B) Quantification of dendritic spine density in CA1 pyramidal neurons shows no significant difference between WT and KO mice. (C) LTP induced by theta-burst stimulation (TBS-LTP) is significantly reduced in KO mice compared to WT controls. (D) Input–output (I/O) relationship showing representative fEPSP slope plotted against afferent fiber volley amplitude for WT and KO slices. (E) Quantification of linear fit slope values from I/O curves shows no significant difference between WT and KO groups. (WT; n = 12 slices, 7 animals, KO; n = 11 slices, 8 animals, Mann-Whitney test, * p < 0.05).

In contrast, activity-dependent synaptic plasticity was significantly impaired. Long-term potentiation (LTP) induced by theta-burst stimulation was markedly reduced in KO mice compared to WT controls (**Fig. 5C**), indicating a deficit in synaptic strengthening mechanisms. Together, these findings demonstrate that *D5Ertd579e* is not required for baseline synaptic structure or transmission but is required for normal synaptic plasticity, revealing a selective functional impairment at the level of activity-dependent circuit modification.

## Discussion

In this study, we identify KIAA0232 (D5Ertd579e) as a previously uncharacterized regulator of neurodevelopment whose disruption results in selective behavioral and synaptic phenotypes without overt structural abnormalities. Despite preserved cortical progenitor organization, laminar patterning, and overall brain morphology, *D5Ertd579e* KO mice exhibited robust impairments in vocalization, sociability, and novelty preference, alongside increased locomotor and perseverative behaviors. In contrast, hippocampal- and amygdala-dependent learning and memory remained intact. Together, these findings indicate that *D5Ertd579e* loss results in domain-specific functional alterations rather than global neurodevelopmental disruption.

A central feature of this phenotype is its selectivity. Behavioral impairments were largely restricted to social and motivational domains, while cognitive functions such as associative learning and memory were preserved. This dissociation is further supported at the circuit level: inhibitory neuron number and synaptic density in the hippocampus were unchanged, and basal synaptic transmission remained intact, yet activity-dependent synaptic plasticity was significantly impaired. These findings suggest that *D5Ertd579e* does not appear to be required for the establishment of neural circuits, but instead contributes to their functional refinement, particularly in processes underlying experience-dependent plasticity.

The behavioral profile observed in *D5Ertd579e* KO mice aligns with emerging models of ASD that emphasize disruption of specific neural circuits rather than global brain dysfunction. In particular, alterations in social behavior, repetitive actions, and locomotor activity are consistent with dysfunction in corticostriatal and cerebellar networks implicated in motivation, motor control, and social cognition (Fucillo, 2016). While these circuits were not directly interrogated in the present study, the convergence of selective behavioral deficits and preserved hippocampal and amygdala-dependent functions supports a model of regionally selective vulnerability.

From a developmental perspective, the absence of large cortical abnormalities suggests that D5Ertd579e does not regulate early neurogenesis or large-scale structural organization.

Instead, its effects may emerge through more subtle processes such as synaptic maturation, circuit integration, or the timing of developmental transitions. This interpretation is consistent with frameworks emphasizing critical periods and activity-dependent refinement in ASD, where disruptions in synaptic pruning or plasticity can lead to long-term circuit dysfunction without gross anatomical changes (LeBlanc & Fagiolini, 2011; Schafer et al., 2012). Notably, impaired synaptic plasticity has been reported across multiple ASD-associated mouse models and has been linked not only to learning deficits, but also to alterations in social behavior and circuit-level information processing (Bourgeron, 2009). For example, disruption of NMDAR-dependent plasticity in Neuroligin-3 mutant mice impairs social memory encoding, while APP family-deficient mice exhibit both impaired LTP and ASD-like social phenotypes (Steubler et al., 2021). In this context, the observed reduction in LTP in D5Ertd579e knockout mice may reflect altered activity-dependent circuit refinement rather than a generalized impairment in hippocampal learning and memory.

Although *D5Ertd579e* remains poorly characterized, limited existing data are broadly consistent with our findings. In Xenopus tropicalis, knockdown of the KIAA0232 orthologue has been reported to reduce telencephalon size, suggesting a role in forebrain development (Naing, 2022). In addition, annotations from the Mouse Genome Informatics database report altered vocalization and behavioral phenotypes associated with *D5Ertd579e* disruption. However, prior work has not directly examined its role in cortical organization, synaptic function, or behavior in mammalian systems. Our findings therefore provide the first integrated characterization of *D5Ertd579e* function across developmental, behavioral, and physiological domains.

Several limitations should be considered. First, the molecular function of KIAA0232 remains undefined, and future studies are needed to identify its interacting partners and downstream pathways. Sequence homology alignments and structural prediction tools tentatively group KIAA0232 within superfamilies that possess ATP-binding or general nucleotide-binding pockets. This suggests it may possess primitive enzymatic, chaperone, or transport activity driven by nucleotide hydrolysis. Indeed, our NetGO 3.0 protein annotation modeling predicts potential transferase activity for this locus (**Figure S1**). Second, while our behavioral analyses revealed robust phenotypes, expanding future investigations to larger cohorts will be essential to catch subtler behavioral trends or uncover potential sex-specific differences. Third, although electrophysiological findings suggest circuit-selective alterations, direct investigation of corticostriatal and cerebellar networks will be required to define the anatomical and functional substrates underlying the observed behavioral phenotypes. Finally, while our knockout mouse model provides an invaluable in vivo platform for system-level behavioral and physiological analyses, cross-species differences in cortical development and synaptic architecture may limit direct translation of these findings to human neurodevelopment. Human-derived models may therefore provide complementary insight into KIAA0232 function in human cortical development. Future studies using isogenic KIAA0232 knockout human iPSC lines differentiated into 3D forebrain organoids could help address this limitation. Such platforms may better recapitulate human-specific progenitor dynamics, cortical developmental timing, and species-specific cellular vulnerabilities that are difficult to model in rodents. Interrogating KIAA0232-deficient human organoids may further clarify whether the selective synaptic phenotypes observed here reflect conserved mechanisms relevant to human cortical circuit development.

In summary, our study identifies KIAA0232 as a novel contributor to neurodevelopment whose disruption leads to selective impairments in social behavior and synaptic plasticity without widespread structural abnormalities. These findings support a model in which uncharacterized genes can influence specific neural circuits and behavioral domains, providing a potential mechanism for how diverse genetic perturbations converge on shared ASD-related phenotypes. More broadly, this work highlights the importance of investigating poorly annotated loci to refine our understanding of neurodevelopmental disorders and their underlying circuit-level mechanisms.

## Author contributions and acknowledgements

D.C. conceived the project and designed experiments. I.S. performed majority of immunostainings, quantifications, and administered behavioral tests. P.J.L. performed electrophysiological experiments and analyzed the data. M.N. performed interneuron stainings and quantifications. Manuscript was written by E.O., D.R., with input from D.C., P.J.L., and M.N. This work was supported by a NIH grant 1R01MH120156-01 to D.C.

