## Supplementary material for "The ASD Risk Gene *D5Ertd579e* Regulates Synaptic Plasticity and Selective Autism-Related Behaviors": Figure S1

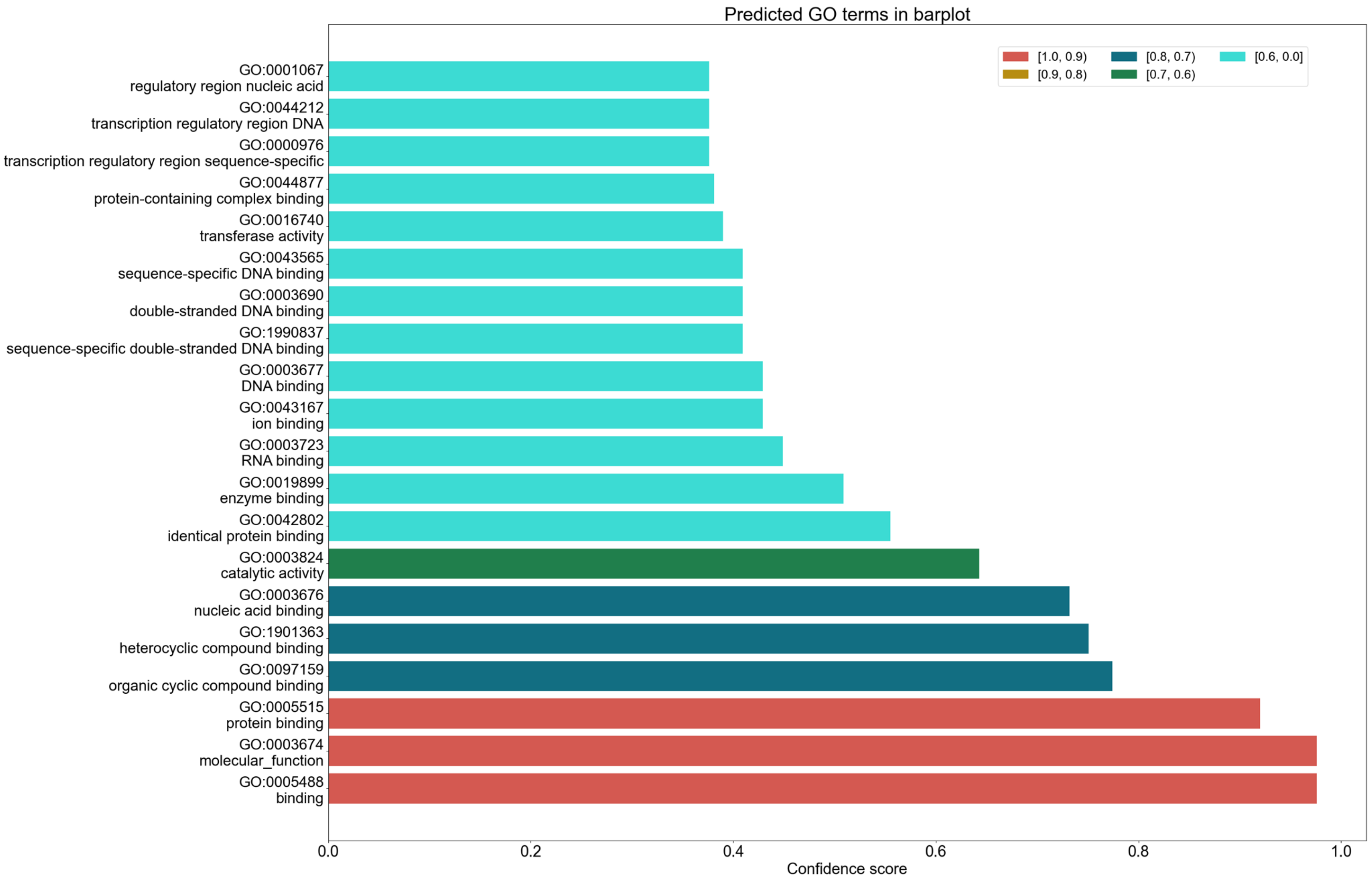
Predicted GO terms in barplot
[1.0, 0.9)
[0.9, 0.8)
[0.8, 0.7)
[0.7, 0.6)
[0.6, 0.0]
GO:0001067
regulatory region nucleic acid
GO:0044212
transcription regulatory region DNA
GO:0000976
transcription regulatory region sequence-specific
GO:0044877
protein-containing complex binding
GO:0016740
transferase activity
GO:0043565
sequence-specific DNA binding
GO:0003690
double-stranded DNA binding
GO:1990837
sequence-specific double-stranded DNA binding
GO:0003677
DNA binding
GO:0043167
ion binding
GO:0003723
RNA binding
GO:0019899
enzyme binding
GO:0042802
identical protein binding
GO:0003824
catalytic activity
GO:0003676
nucleic acid binding
GO:1901363
heterocyclic compound binding
GO:0097159
organic cyclic compound binding
GO:0005515
protein binding
GO:0003674
molecular_function
GO:0005488
binding
0.0
0.2
0.4
0.6
0.8
1.0
Confidence score

**Supplemental Figure 1. Predicted molecular function Gene Ontology (GO) annotations for KIAA0232 generated using NetGO 3.0.**

The full UniProt amino acid sequence of KIAA0232 was analyzed using the NetGO 3.0 protein annotation platform. The bar plot shows the top predicted molecular function GO terms ranked by confidence score. Bar colors indicate confidence score ranges as shown in the legend. Predicted annotations include binding-related functions, nucleic acid binding, catalytic activity, and transferase activity.
